# Predicting Human Bioavailability of Subcutaneously Administered Fusion Proteins and Monoclonal Antibodies

**DOI:** 10.1101/2023.01.15.524112

**Authors:** Peng Zou

## Abstract

There has been an increasing trend towards subcutaneous (SC) delivery of fusion proteins and monoclonal antibodies (mAbs) in recent years versus intravenous (IV) administration. The prediction of bioavailability is one of the major barriers in clinical translation of SC administered therapeutic proteins due to a lack of reliable in vitro and preclinical in vivo predictive models. In this study, we explored the relationships between human SC bioavailability and physicochemical or pharmacokinetic properties of 20 Fc-or albumin-fusion proteins and 98 monoclonal antibodies. An inverse linear correlation was observed between human SC bioavailability and human intravenous clearance (CL) or isoelectric point (pI). The bioavailability of fusion proteins is more correlated with pI while the bioavailability of mAbs is more correlated with CL. A mAbs with intravenous CL < 4 mL/day/kg is likely to have SC bioavailability > 60%. Multivariate regression models were developed using intravenous CL and pI of a training set (N = 59) as independent variables. The predictive models were validated with an independent test set (N = 33). A linear regression model resulted in 27 among 33 (82%) predictions within 0.8-to 1.2-fold deviations. Overall, this study demonstrated that CL- and pI-based multivariate regression models could be used to predict human SC bioavailability of fusion proteins and mAbs.

## 1. Introduction

According to a recent industry-wide survey, the most frequently used routes of administration of therapeutic proteins are intravenous (68%), followed by subcutaneous (23%), and others (e.g., intravitreal and intratumoral, 9%)[1]. Subcutaneous (SC) drug products can be self-administered by a patient or administered by another person at home, which is an advantage over intravenous (IV) administration especially for chronic use. There has been an increasing trend towards SC delivery of fusion proteins and monoclonal antibodies (mAbs) in recent years versus IV administration. Despite advantages of SC administration of therapeutic proteins, there remain critical development issues and knowledge gaps in SC drug delivery that need to be addressed to further progress the field. To address the knowledge gaps in SC delivery of therapeutic proteins, the SC Drug Delivery and Development Consortium was convened in 2018 as a collaboration of well recognized industry experts. In the article published by the Consortium in 2020[2], SC bioavailability prediction is identified as one of the key knowledge gaps in SC delivery of therapeutic proteins, with a high priority second to the development of high dose/volume SC formulations. Currently, there are no reliable preclinical models available to accurately predict the bioavailability of SC administered proteins across species[2]. In 2020, the Consortium issued an open challenge to industry and academia, encouraging the development of reliable models to enable SC bioavailability prediction of therapeutic proteins in humans and improve translation from preclinical species [3].

An in vitro diffusion model termed Scissor was developed to predict human SC bioavailability of eight mAbs [4]. Although the in vitro model exhibited a higher prediction accuracy than monkey and minipig models, the performance of the model has not been verified with a large number of mAbs. An equation (F% = -6.72 * CL + 89.4) was generated from a linear regression between observed human SC bioavailability and clearance of 19 mAbs and used to predict human SC bioavailability of mAbs[5]. However, the validity of the equation was not evaluated with an independent dataset. Recently, machine learning algorithms were developed by fitting 47 features of 36 mAbs in a training set [6]. Model validation with 9 mAbs showed the machine learning algorithms could predict whether a mAb has a SC bioavailability ≥70% or not but could not provide a quantitative prediction.

During past several years, more and more human SC bioavailability data of fusion proteins and mAbs have been published. This author believes it is an appropriate time to summarize human SC bioavailability data, investigate the key factors affecting human SC bioavailability, and fill the knowledge gap in human SC bioavailability prediction. The physicochemical properties of protein drugs (electric charges, hydrophobicity, FcRn binding affinity, catabolic stability, hydrophobicity, molecular weight et al.), intrinsic factors (human obesity, immunogenicity, lymphatic flow rate, injection site et al.) and extrinsic factors (dose, formulation composition, injection volume, viscosity) that influence the absorption of fusion proteins and mAbs following SC administration have been discussed previously [7-10]. Among these factors, the effect of electrostatic and hydrophobic interaction between protein drugs and SC tissue on the SC bioavailability of six humanized mAbs was evaluated in both rats and cynomolgus monkeys [11]. The study showed that a combination of high positive charge and hydrophobic interaction significantly reduced the rate of absorption and bioavailability of the six mAbs. At physiological pH, the interstitial space of SC tissue is overall negatively charged because hyaluronic acid, chondroitin sulfate and proteoglycans are negatively charged, resulting in faster transport of negatively charged biologics compared to positively charged ones [10]. Furthermore, it was reported that mAbs exhibited incrementally enhanced binding to cell surfaces and cellular uptake with increased positive charge in antigen-negative cells [12]. Thus, compared to negatively charged and neutral mAbs, positively charged mAbs may show enhanced binding to epithelial cells and cellular uptake in SC tissue. The longer SC retention of positively charged mAbs may render them to increased extracellular catabolism.

SC presystemic catabolism through extracellular proteolysis and macrophage or dendritic cell clearance may reduce the bioavailability of protein drugs[3]. The proteolysis and macrophage/dendritic cell uptake occur in both SC extracellular space and during lymphatic transport. FcRn binding may facilitate the recycling of mAbs and Fc or albumin-fusion proteins and protect them from degradation in macrophages and dendritic cells [3]. Therefore, catabolic stability and FcRn binding affinity are important factors which affect SC bioavailability of mAbs and fusion proteins. Unfortunately, in vitro catabolic stability of most mAbs and fusion proteins are not available in literature. Furthermore, a large inter-lab variability in FcRn binding affinity measurements precludes a meaningful correlation analysis between SC bioavailability of mAbs or Fc-fusion proteins and in vitro FcRn binding affinity reported in literature. An in vitro FcRn-dependent transcytosis assay was successfully developed to predict the linear clearance of 53 mAbs, indicating a correlation between nonspecific linear clearance of mAbs and FcRn binding[13]. Furthermore, a correlation analysis showed an inverse linear relationship between human SC bioavailability and clearance of 19 mAbs [5]. Human clearance of mAbs and fusion proteins may be predictive of human SC bioavailability.

Similar to the application of Biopharmaceutics Classification System (BCS) to the oral bioavailability prediction for small-molecule drugs, the SC Drug Delivery and Development Consortium proposed that a drug classification system might be developed for SC administered mAbs. To respond the Consortium’s open challenge, in current study, the author collected human SC bioavailability data of fusion proteins and mAbs currently available to public and assessed the potential relationships between SC bioavailability and physicochemical properties/pharmacokinetic parameters of these therapeutic proteins. Recently, Ahmed et. al. calculated 24 physicochemical descriptors of variable fragment (Fv) regions for 77 marketed mAbs and 271 mAbs under clinical development using homology-based molecular models[14]. Among the 24 descriptors, five descriptors (the surface area buried between the variable domains of light chain and heavy chain, 3D structure-based isoelectric point of Fv region, ratio of dipole and hydrophobic moments, ratio surface area of charged to hydrophobic patches, and average hydrophobic imbalance) were identified as nonredundant descriptors because no statistically significant correlations were observed among them. The five descriptors were found to be able to describe the stability, electrostatics, hydrophobicity, and molecular surface properties of Fv regions [14]. Therefore, all the five descriptors of Fv region were included in the correlation analysis with SC bioavailability in current study.

Based on the correlations observed between human SC bioavailability and physicochemical properties/pharmacokinetic parameters, the author proposed a SC bioavailability classification system and developed multivariate regression models to predict human SC bioavailability of fusion proteins and mAbs.

## 2. Methods

### 2.1 Data source

Only standard mAbs and Fc-or albumin-fusion proteins were collected. Proteins with modified structures (e.g., scFv, Fab fragment, antibody-drug conjugates, pegylated mAb) were not taken into consideration in present study. For SC drug products approved by the FDA, product labels and clinical pharmacology reviews were surveyed to collect human PK data. European public assessment reports database (https://www.ema.europa.eu/en/medicines) was searched to identify mAbs and fusion proteins. The list of therapeutic antibodies from the antibody society website (https://www.antibodysociety.org/resources/approved-antibodies/) was also used to identify SC administered antibodies. In addition to the FDA-or EMA-approved or reviewed mAbs and fusion proteins, the author collected mAbs and fusion proteins at clinical development stage from the Antibodies to Watch published by mAbs Journal between 2010-2022 [15-19] and three research articles [14, 20, 21]. A total of more than 500 mAbs and fusion proteins approved by the FDA/EMA or under clinical development was collected. Then, a thorough literature search was conducted in PubMed and Google to collect the SC bioavailability, linear clearance and nonlinear clearance following an IV dose of these proteins. When different human SC bioavailability values at various dose levels were available, the bioavailability at a high dose or from a cohort of large sample size was selected. When only a range of F was reported, the mean value is selected. When available, the intravenous clearance determined at the dose used for absolute bioavailability study was selected. If available, both linear clearance (at a high dose) and nonlinear clearance (at the lowest dose) following IV administration were collected separately.

When only human plasma concentration-time curves were available, an online graph reader tool (http://www.graphreader.com/) was used to digitalize the PK data for PK parameter calculation.

Five physicochemical descriptors of mAbs including the surface area buried between the variable domains of light chain and heavy chain (BSA_VL:VH), 3D structure-based isoelectric point of Fv region (3D_pI), ratio of dipole and hydrophobic moments (RM), ratio of surface area of charged to hydrophobic patches (RP), and average hydrophobic imbalance (Avg_HI) were collected from a literature report[14]. The amino acid sequences of mAbs and fusion proteins were collected from NCATS Inxight Drugs (https://drugs.ncats.io/). The sequence-based isoelectric point (pI) of the whole protein molecule was calculated based on the amino acid sequence using an online tool in Expasy (https://web.expasy.org/compute_pi/).

### 2.2 Correlation analysis and drug classification

Linear regression analysis was conducted between SC bioavailability and each individual physicochemical property or clearance. The analysis was conducted for fusion proteins and mAbs separately. The two factors showing a higher correlation coefficient were used to classify mAbs into four subclasses.

### 2.3 Multivariate regression

The multivariate regression analysis was conducted for fusion proteins and mAbs separately. The two physicochemical or PK properties showing a correlation with SC bioavailability were selected as the independent variables for multivariate regression in SigmaPlot (version 14.0). Plane and paraboloid fittings were used to generate the regression equations. All fusion proteins were used in multivariate regression. The mAbs with name starting with letter A to N were assigned to a training set and the other mAbs were assigned to a test set. Only the training set was used for multivariate regression.

### 2.4. Prediction accuracy of regression models

The regression models were assessed for accuracy using the test set. The percentage of predictions within 0.8-to 1.2-fold error (PPwFE) and geometric mean fold error (GMFE) were the metrics to evaluate the performance of different regression models. PPwFE and GMEFE were calculated by eq 1 and eq 2, respectively.

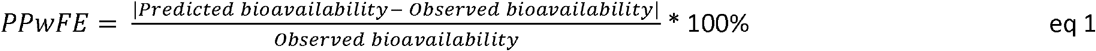

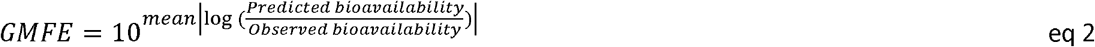

## 3. Results

### 3.1 Overview of the dataset

A total of 20 fusion proteins (19 Fc-fusion proteins and 1 albumin-fusion protein) and 98 mAbs with human SC bioavailability available were collected. The five physicochemical descriptors (BSA_VL:VH, 3D_pI, RM, RP, and Avg_HI) of fusion proteins were not available. The sequence-based pI and IV linear clearance of the 20 fusion proteins were available (Table S1). Among the 98 mAbs (Table S2), the five in silico physicochemical descriptors (BSA_VL:VH, 3D_pI, RM, RP, and Avg_HI) of 79 mAbs were available in literature report [14]. The amino acid sequences of 2 mAbs (CNTO 5825 and PF-04236921) were unknown. Thus, the sequence-based pI was calculated for 96 mAbs. The linear clearance of 4 mAbs (brazikumab, lirentelimab, pozelimab, and rosnilimab) following an IV dose was not available. Finally, 92 mAbs with SC bioavailability, sequence-based pI, and linear IV clearance available were assigned to training and test sets for multivariate regression analysis. A total of 59 mAbs with name starting with A to N was assigned to training set and the remaining 33 mAbs were assigned to test set. The nonlinear clearance (target-mediated drug disposition clearance at a low dose) was available for 45 mAbs (Table S2).

### 3.2 Regression analysis for fusion proteins

A linear correlation analysis was conducted for 20 fusion proteins between human SC bioavailability and sequence-based pI or IV linear clearance. The 20 fusion proteins and their SC bioavailability, whole molecule sequence-based pI, and IV linear clearance were summarized in Supporting Materials Table S1. A moderate inverse correlation was observed between SC bioavailability and sequence-based pI (Figure 1A) or IV linear clearance (Figure 1B), indicating SC electric interaction and catabolism affect SC bioavailability of fusion proteins. Then, sequence-based pI and IV linear clearance were used as independent variables for multivariate regression analysis (Figure 2). A linear equation (F = 1.2201 – 0.0955 × pI – 0.0012 × CL) was derived and the regression coefficient (R^2^) was 0.541 (Table 1). Due to the small number of fusion protein (N = 20), model validation with an independent test set was not conducted.

**Figure 1.**
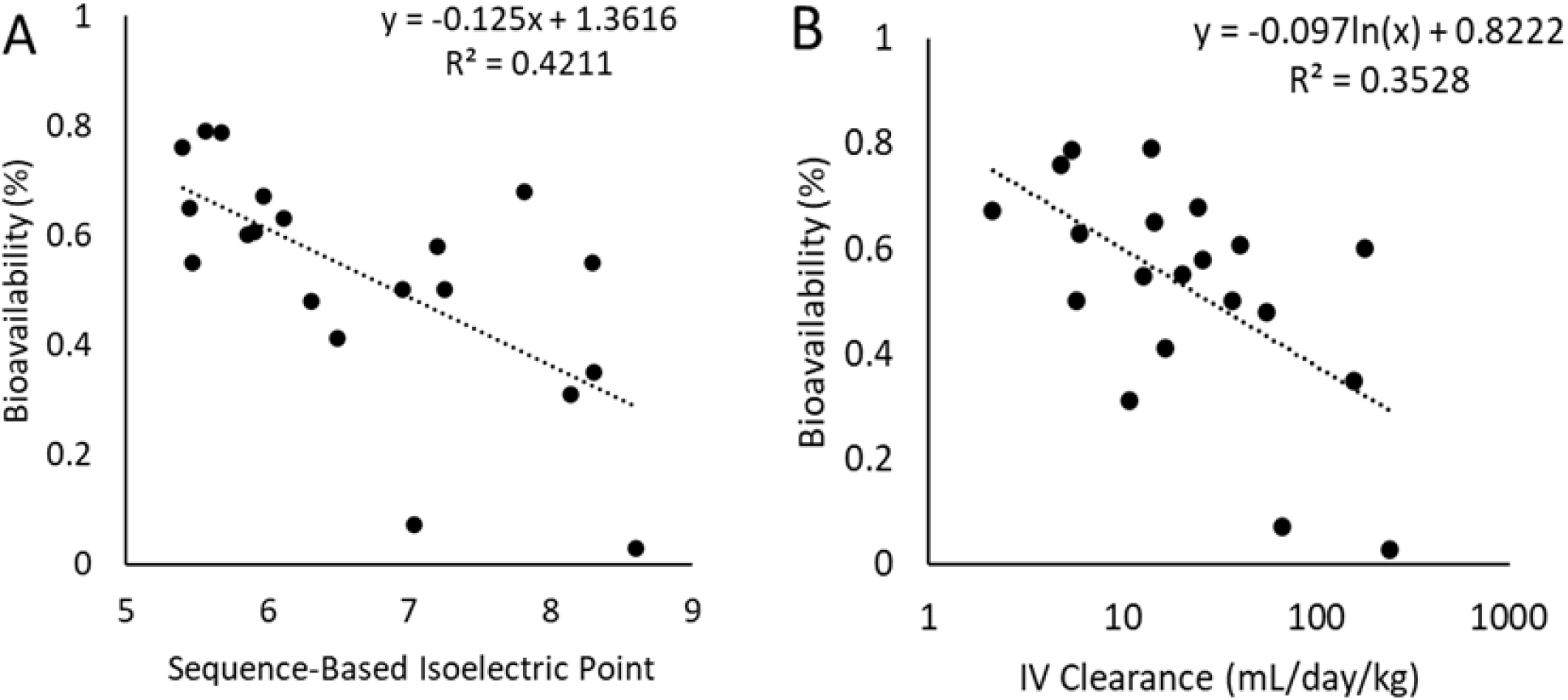
Relationships between human subcutaneous bioavailability of fusion proteins and (A) sequence-based isoelectric point or (B) intravenous linear clearance (N = 20).

**Figure 2.**
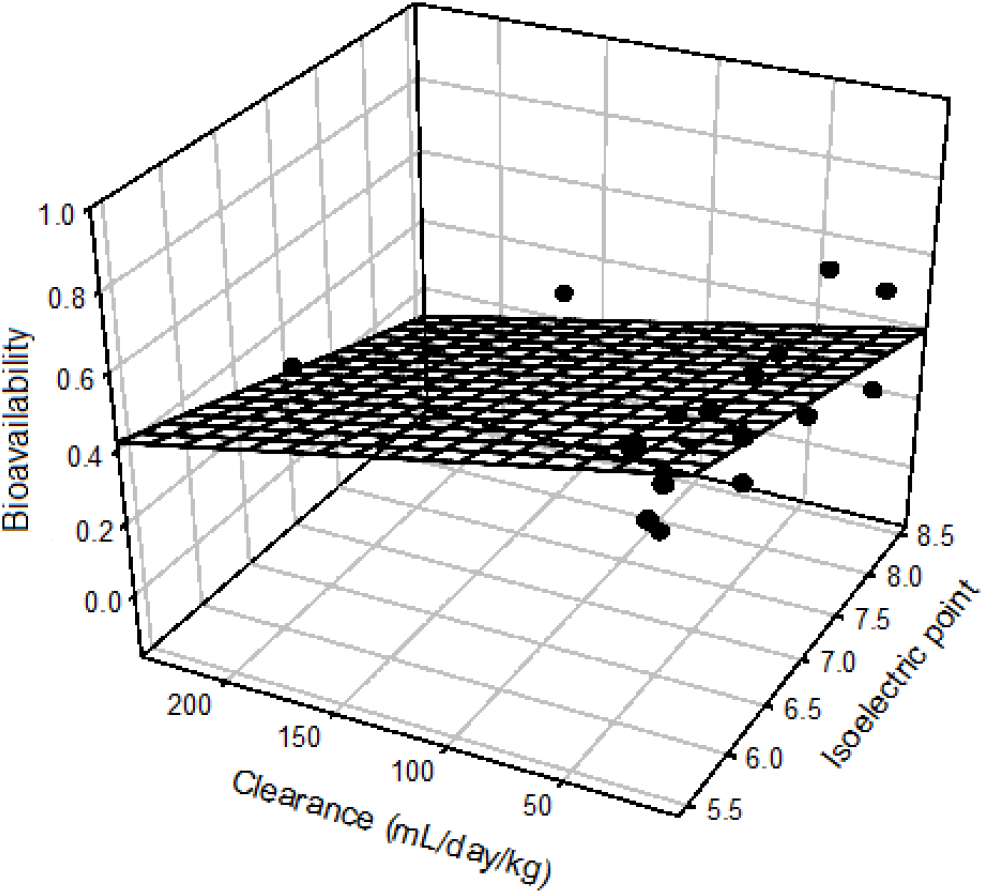
Linear multivariate correlation of subcutaneous bioavailability for fusion proteins (N = 20)

**Table 1.**
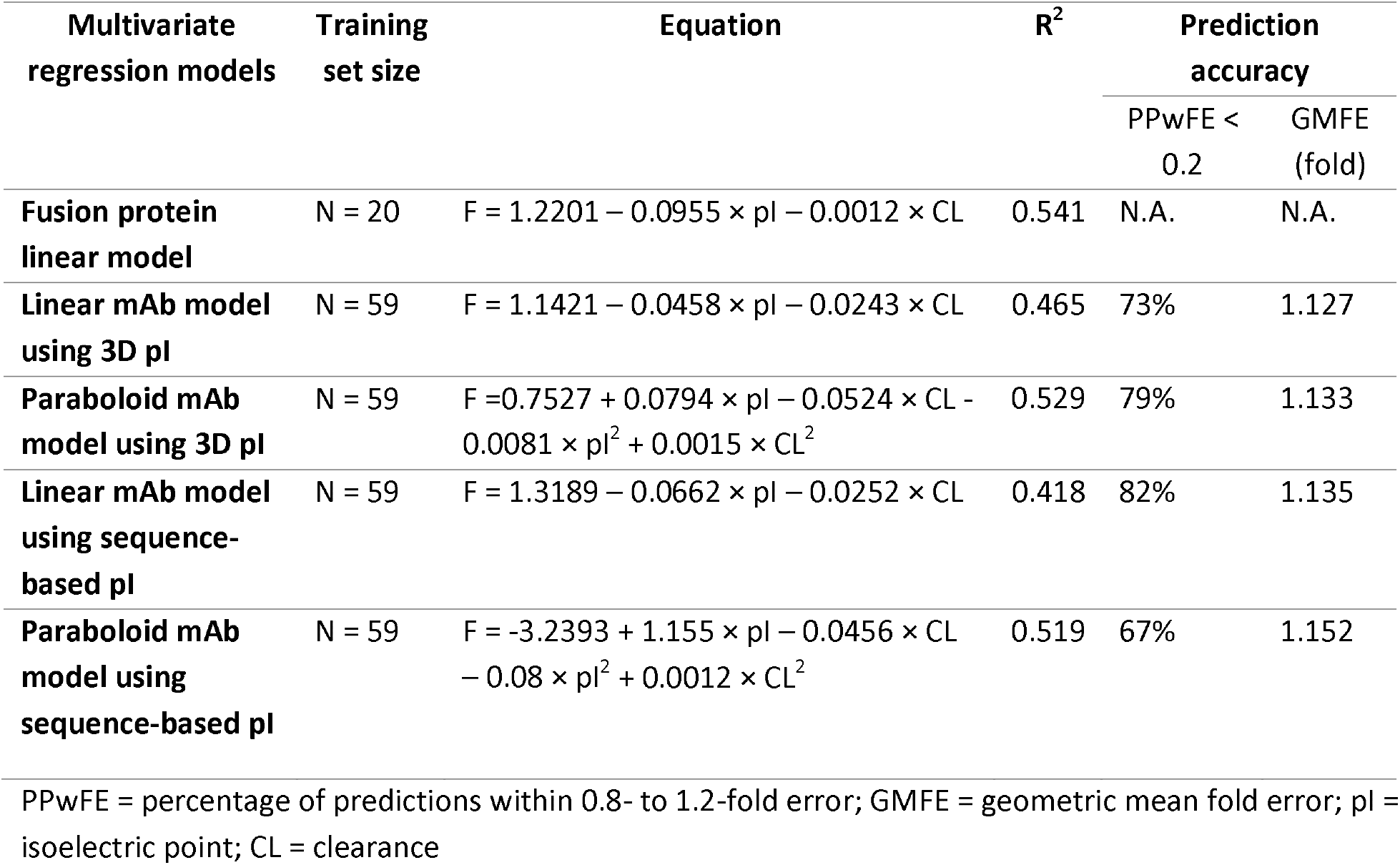
Comparison of the performance of multivariate regression models for fusion proteins and mAbs.

### 3.3 Correlation analysis and drug classification for mAbs

Six physicochemical parameters (sequence-based pI, 3D_pI, BSA_VL:VH, RM, RP, and Avg_HI) and two pharmacokinetic parameters (linear clearance and nonlinear clearance) of were tested in correlation analysis with SC bioavailability of mAbs. The physicochemical and pharmacokinetic parameters and SC bioavailability of 98 mAbs were summarized in Supporting Materials Table S2. No correlation was observed between SC bioavailability and BSA_VL:VH, RM, RP, or Avg_HI (R^2^ < 0.02) (Figure 3). In contrast, a week correlation was observed between SC bioavailability of mAbs and 3D_pI (R^2^ = 0.174) or sequence-based pI (R^2^ = 0.092). Consistent with the correlation observed with fusion proteins, a moderate correlation was observed between SC bioavailability of mAbs and linear IV clearance (R ^2^= 0.343) or nonlinear clearance (R^2^ = 0.250) (Figure 4).

**Figure 3.**
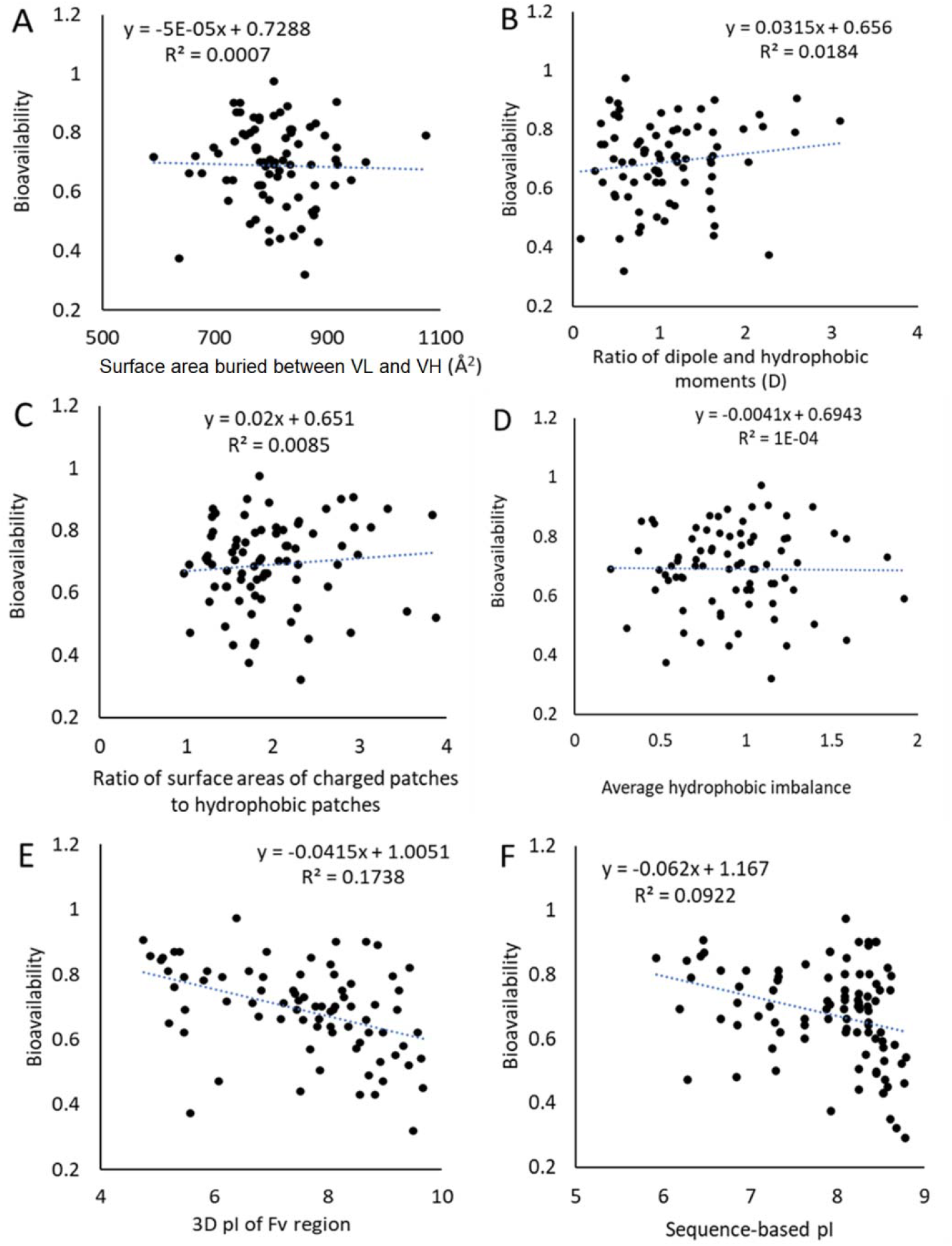
Relationships between human subcutaneous bioavailability of mAbs and physicochemical parameters. (A) surface area buried between variable domains of light chain and heavy chain (N = 79), (B) ratio of dipole and hydrophobic moments (N = 79), (C) ratio of surface area of charged to hydrophobic patches (N = 79), (D) average hydrophobic imbalance (N = 79), (E) 3D structure-based isoelectric point of Fv region (N = 79), and (F) sequence-based isoelectric point of mAb (N = 96).

**Figure 4.**
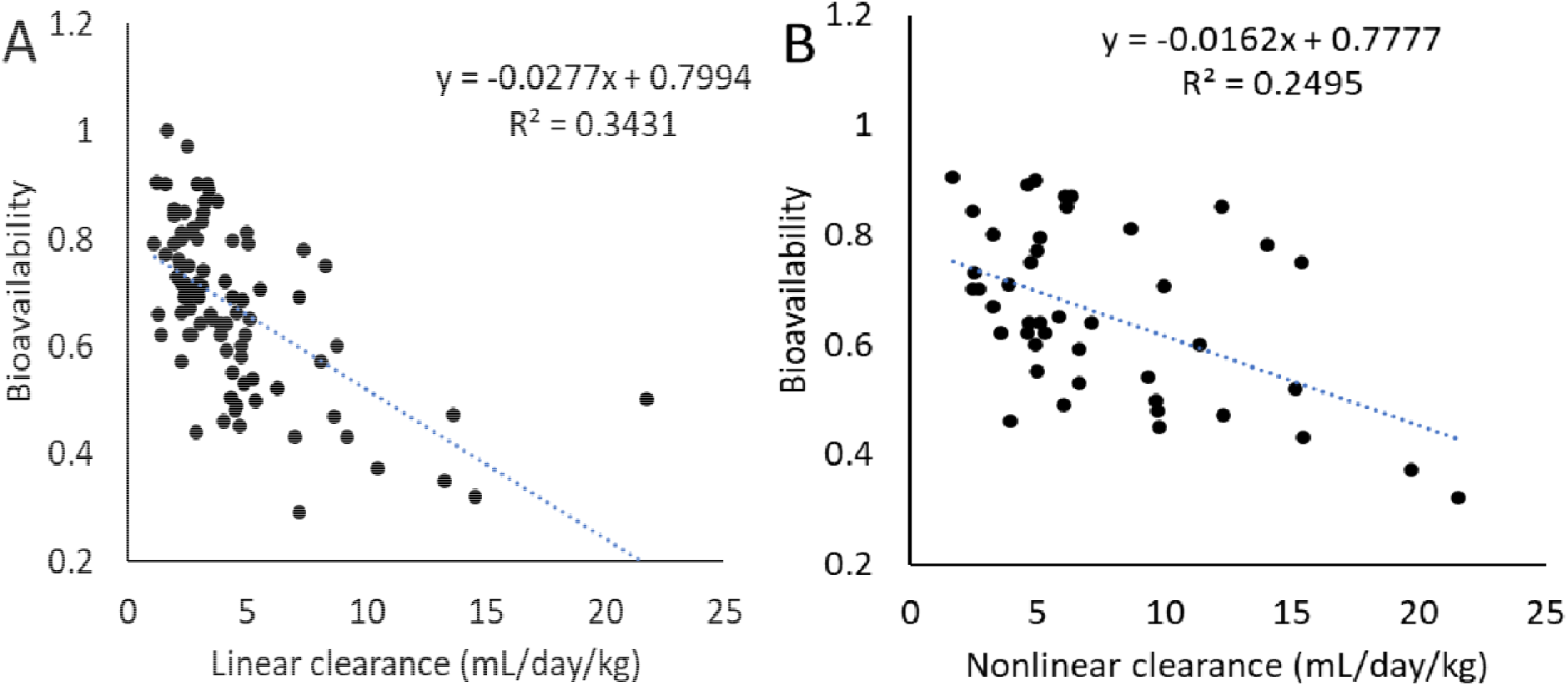
Relationships between human subcutaneous bioavailability of mAbs and intravenous pharmacokinetic parameters. (A) linear clearance (N = 94) and (B) nonlinear clearance (N = 45)

Based on the correlations observed between SC bioavailability and IV linear clearance/3D pI of Fv region, the two parameters were used to classify mAbs. As shown in Table S2, 79 mAbs had both 3D pI of Fv region and sequence-based pI available while 17 mAbs had sequence-based pI available only. For the 79 mAbs, the 3D pI of Fv region and sequence-based pI were overall comparable. Therefore, to extend 3D pI-based mAb classification to 96 mAbs, the author assumed the 3D pI of Fv region of the 17 mAbs was the same as their sequence-based pI. The 96 mAbs were assigned to four classes: Class 1, linear clearance < 4 mL/day/kg and 3D pI < 7.5 (N = 23); Class 2, linear clearance < 4 mL/day/kg and 3D pI ≥ 7.5 (N = 31); Class 3, linear clearance ≥ 4 mL/day/kg and 3D pI < 7.5 (N = 9); and Class 4, linear clearance ≥ 4 mL/day/kg and 3D pI ≥ 7.5 (N = 29). Compared to 3D pI of 7.5, linear clearance of 4 mL/day/kg was more effective in distinguishing mAbs with low and high SC bioavailability. Among 19 mAbs with SC bioavailability > 80%, 18 had IV linear clearance < 4 mL/day/kg (Figure 5A). While among 15 mAbs with SC bioavailability ≤ 50%, 14 had IV linear clearance ≥ 4 mL/day/kg. All 23 Class 1 mAbs had SC bioavailability > 60% and 52 among 54 (96%) Classes 1 and 2 mAbs (clearance < 4 mL/day/kg) had SC bioavailability > 60% (Figure 5B).

**Figure 5.**
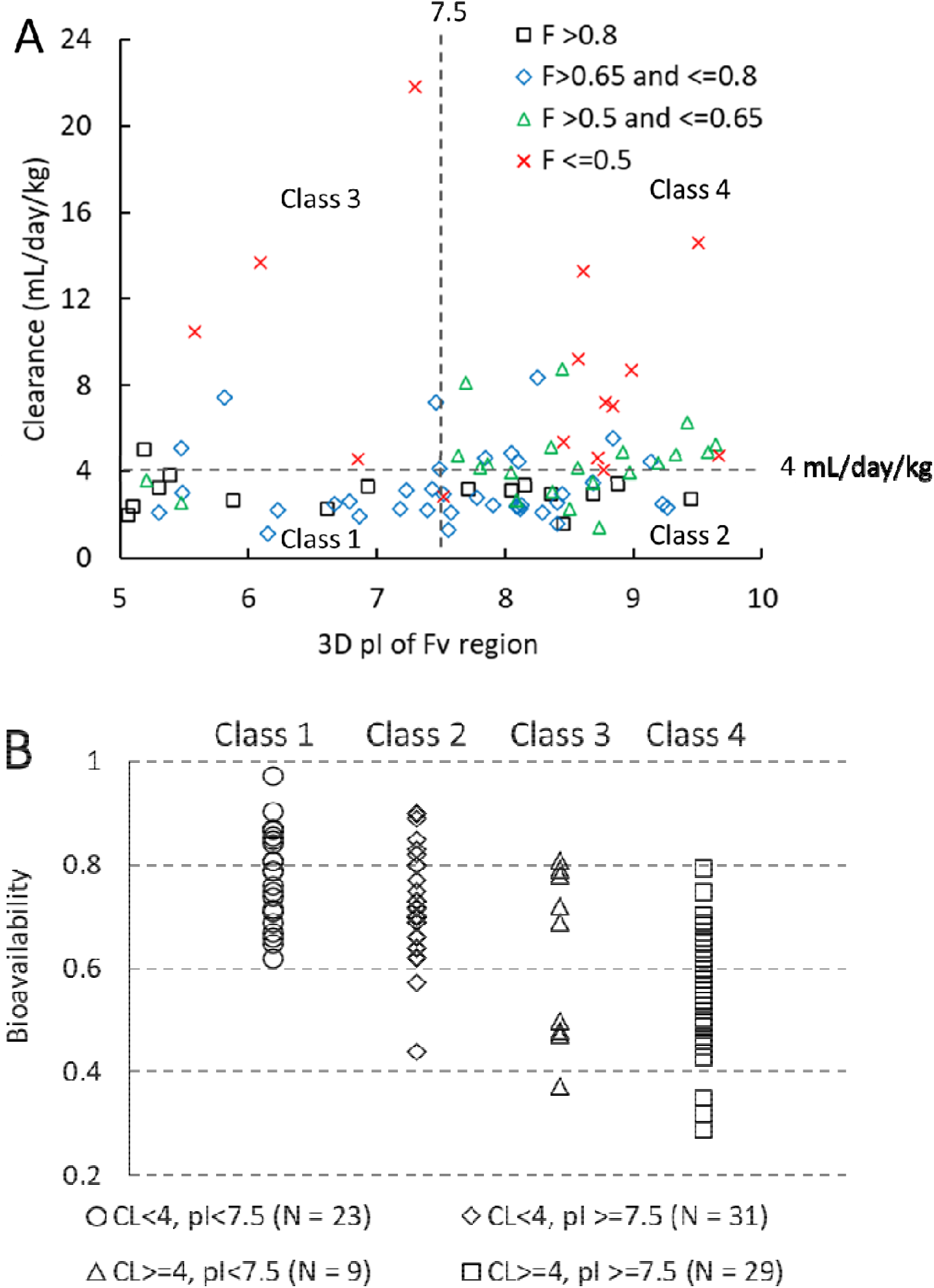
Distribution of subcutaneous bioavailability of 92 mAbs in four classes.

### 3.4 Multivariate regression analysis for mAbs

As shown in Figure 5, the proposed mAb classification system was unable to accurately the SC bioavailability of mAbs. Thus, multivariate regression was explored for SC bioavailability prediction. The linear clearance and isoelectric point were selected as the independent variables for multivariate regression analysis. Both 3D pI of Fv region and sequence-based pI were assessed in the multivariate regression analysis. No obvious correlation was observed between IV linear clearance and 3D pI of Fv region or sequence-based pI (Supporting Materials, Figure S1). The linear and paraboloid regression fitting with IV linear clearance and 3D pI of Fv region of 59 mAbs in the training set resulted in R values of 0.465 and 0.529, respectively (Figure 6). The prediction performance of the linear and paraboloid models was evaluated using an independent test set of 33 mAbs (Figure 7). The linear and paraboloid models resulted in 73% (24/33) and 79% (26/33) of predictions within 0.8-to 1.2-fold errors. The GMFEs of SC bioavailability predicted by linear and paraboloid models were 1.127-fold and 1.133-fold, respectively (Table 1).

**Figure 6.**
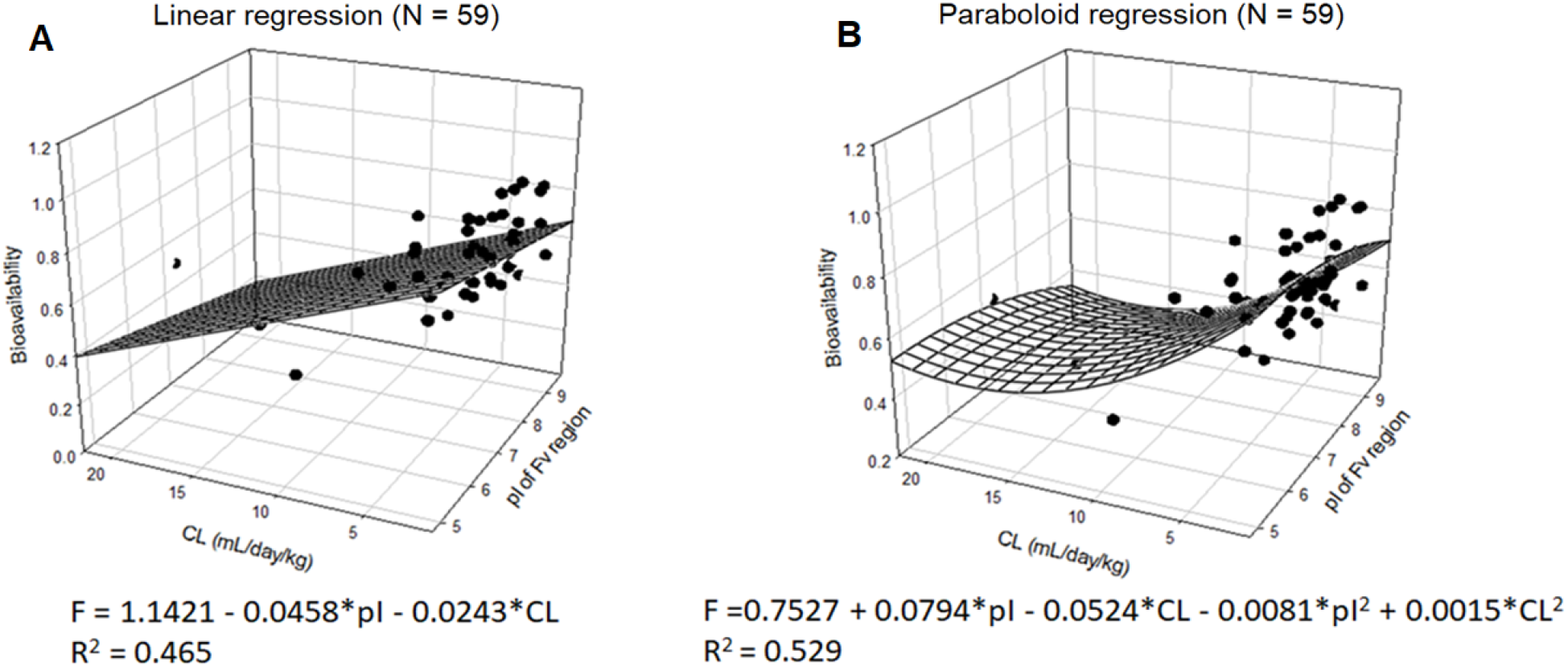
(A) Linear and (B) paraboloid multivariate regression of subcutaneous bioavailability with linear clearance and 3D pI of Fv region (N = 59)

**Figure 7.**
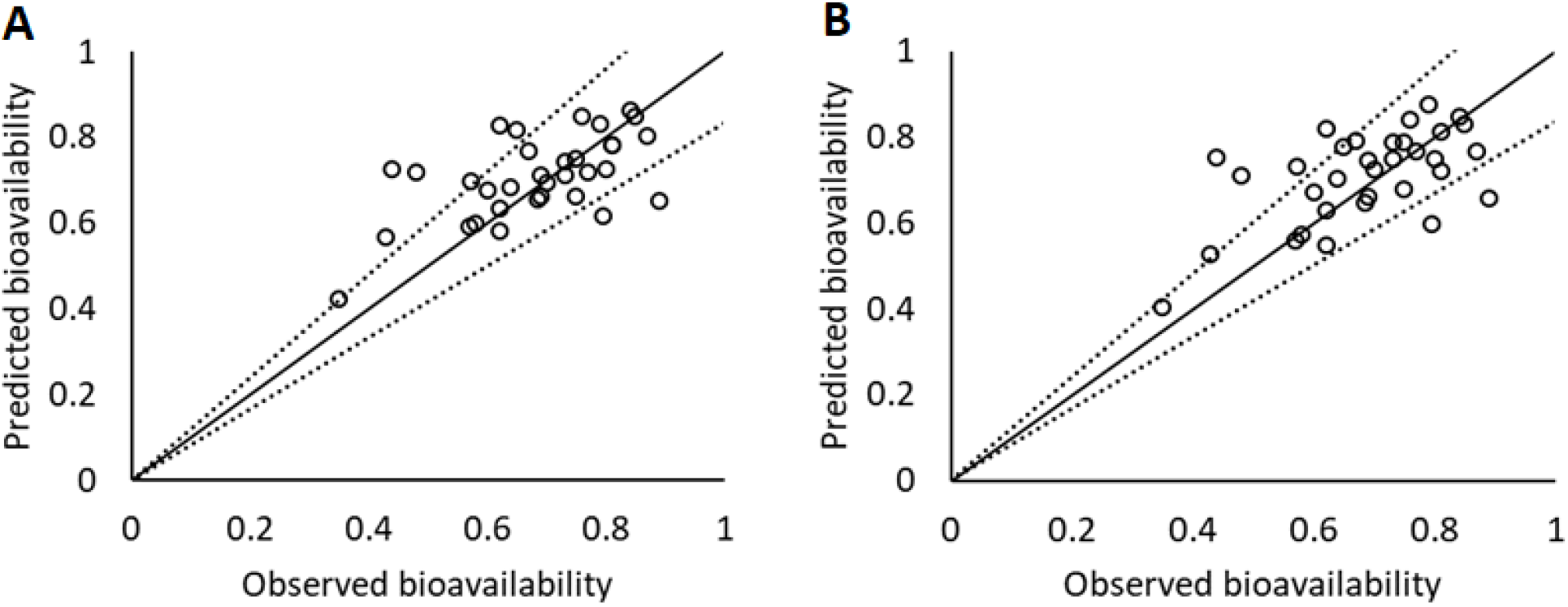
Prediction performance of (A) linear and (B) paraboloid multivariate regression models using 3D pI of Fv region (N = 33). Dash lines are 1.2-fold and 0.8-fold error lines. Solid line represents unity.

The 3D pI of Fv region was calculated by homology modeling using commercial software molecular operating environment (MOE). In contrast, it is more convenient to calculate sequence-based pI of mAbs using an online tool in Expasy. Therefore, sequence-based pI of mAbs was also used for multivariate regression analysis. The linear and paraboloid regression fitting with IV linear clearance and sequence-based pI of 59 mAbs in training set resulted in R^2^ values of 0.418 and 0.519, respectively (Figure 8). The linear and paraboloid models were applied to the 33 mAbs in test set and resulted in 82% (27/33) and 67% (22/33) of predictions within 0.8-to 1.2-fold errors (Figure 9). The GMFEs of SC bioavailability predicted by linear and paraboloid models were 1.135-fold and 1.152-fold, respectively (Table 1).

**Figure 8.**
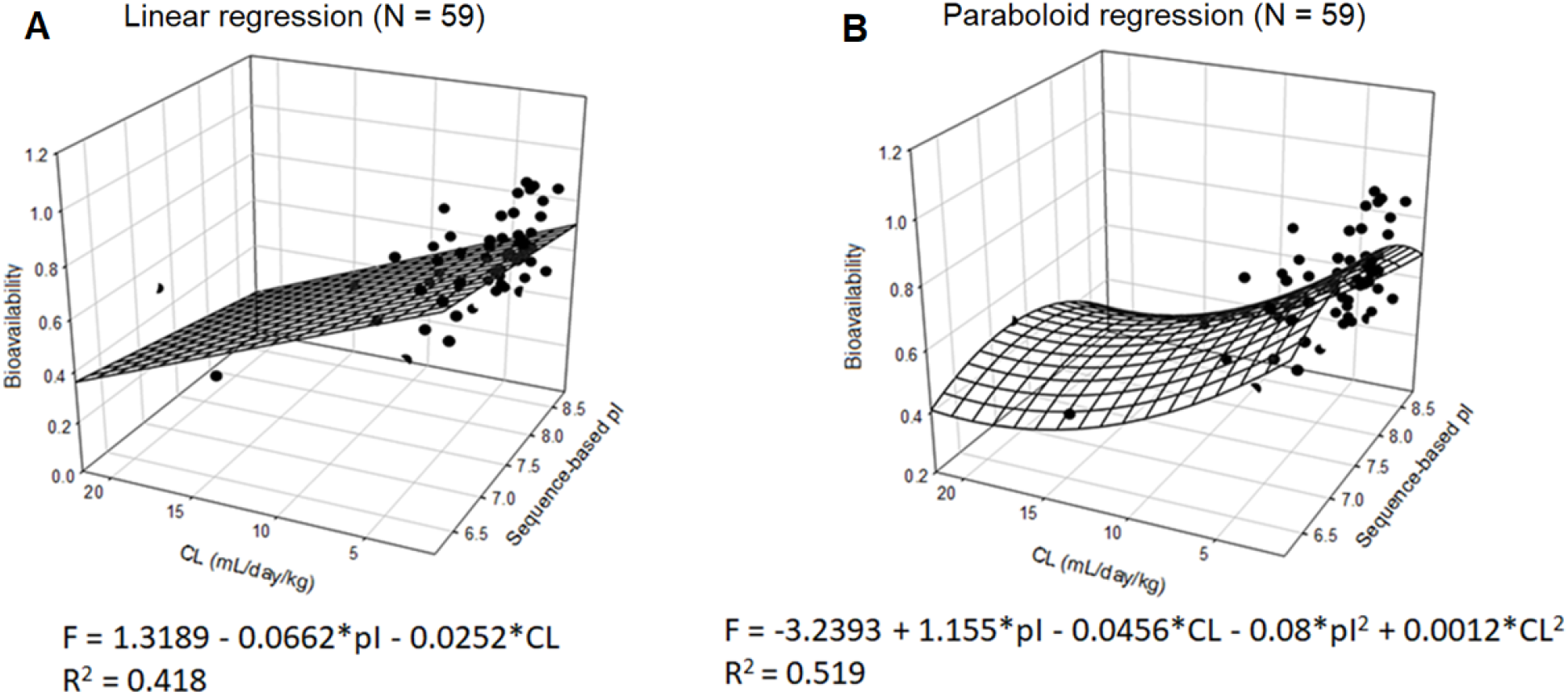
(A) Linear and (B) paraboloid multivariate regression of subcutaneous bioavailability with linear clearance and sequence-based pI (N = 59)

**Figure 9.**
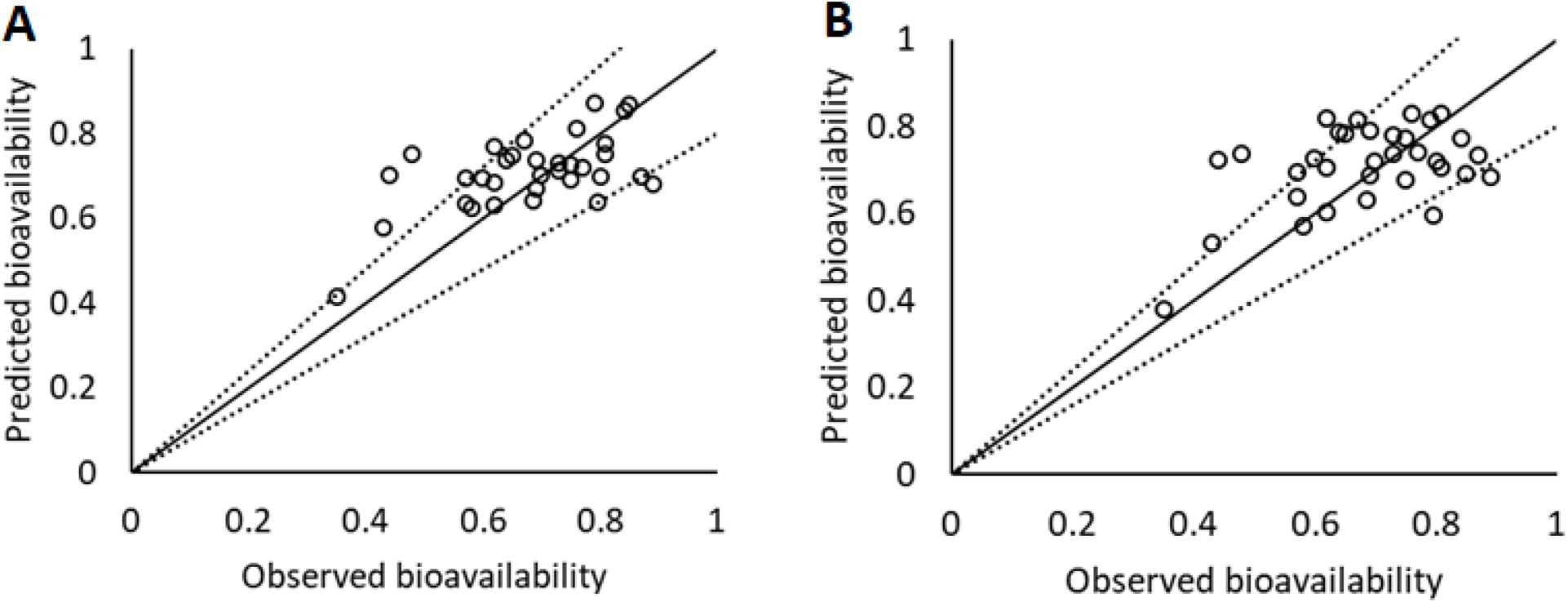
Prediction performance of (A) linear and (B) paraboloid multivariate regression models using sequence-based pI (N = 33). Dash lines are 1.2-fold and 0.8-fold error lines. Solid line represents unity.

## 4. Discussion

In current study, an inverse linear correlation was observed between human SC bioavailability and human intravenous clearance or pI using the data of 20 fusion proteins and 98 mAbs. In contrast, no correlation was observed between SC bioavailability and four hydrophobic properties (the surface area buried between the variable domains of light chain and heavy chain, ratio of dipole and hydrophobic moments, ratio surface area of charged to hydrophobic patches, and average hydrophobic imbalance). The SC bioavailability of fusion proteins is more correlated with pI than linear clearance (R^2^ values for pI and linear clearance are 0.42 and 0.35, respectively) while the SC bioavailability of mAbs is more correlated with linear clearance than 3D pI of Fv region (R^2^ values for 3D pI of Fv region and linear clearance are 0.17 and 0.34, respectively). The higher sensitivity of the SC bioavailability of fusion proteins to pI is likely due to their vulnerability to pre-systemic catabolism. The mean linear clearance of 20 fusion proteins is 47.2 mL/day/kg (Table S1) while the mean linear clearance of 94 mAbs is 4.31 mL/day/kg (Table S1). The average linear clearance of fusion proteins is averagely 11-fold of that of mAbs. Because the interstitial space of SC tissue is overall negatively charged at physiological pH, positive charge (pI > 7.4) is likely to increase the retention time of SC administered proteins. The vulnerability of fusion proteins to pre-systemic catabolism may cause the SC bioavailability of fusion proteins to be sensitive to the changes in pI.

The proposed drug classification based on the linear clearance of mAbs and 3D pI of Fv region showed that mAbs with a linear clearance < 4 mL/day/kg is likely to have SC bioavailability > 60%. However, the proposed cut-off values of linear clearance (4 mL/day/kg) and pI (7.5) were unable to accurately distinguish mAbs with low, moderate, and high SC bioavailability (Figure 5). It is speculated that there are other unidentified physicochemical properties and extrinsic factors (i.e., dose, formulation, viscosity, injection volume and others) which substantially affect SC bioavailability of fusion proteins and mAbs. Further investigation is warranted to identify the relationships between SC bioavailability and other important factors.

It has been well known that pre-systemic catabolism and electric interactions with SC tissues may affect SC absorption of therapeutic proteins. The potential relationship between SC bioavailability and clearance of mAbs was previously assessed using clinical or preclinical data in several reports. For example, a correlation analysis of 8 mAbs showed that minipig clearance was predictive of human linear clearance (coefficient of determination R^2^ = 0.69) and the SC bioavailability appeared to inversely correlate with systemic clearance in minipigs (R^2^ = 0.82), although the SC bioavailability values in minipigs were poorly correlated with those in humans[22]. Similarly, a linear inverse correlation (regression equation F% = - 6.72 * CL + 89.4, R^2^ = 0.62) was observed between SC bioavailability and clearance of 19 mAbs in humans[5]. Our regression analysis of 20 fusion proteins and 94 mAbs further confirmed a moderate linear inverse correlation between SC bioavailability and IV linear clearance of fusion proteins (R^2^ = 0.35) and mAbs (R^2^ = 0.34) in humans. The proteins in human SC interstitial space are qualitatively similar to those in human plasma although their concentrations are approximately 50% lower than that in plasma[23]. Therefore, fusion proteins and mAbs vulnerable to proteolysis in human plasma following IV administration are expected to undergo similar presystemic catabolism in SC interstitial space and lymph fluid[10]. This may explain the moderate inverse correlation between SC bioavailability and IV linear clearance of fusion proteins and mAbs. A linear inverse but weaker correlation (R ^2^= 0.25) was also observed between SC bioavailability and IV nonlinear clearance of 45 mAbs in humans. The weaker correlation with nonlinear clearance may be explained that targeted-mediated elimination of fusion proteins and mAbs in SC tissues is minimal and non-specific presystemic catabolism is dominant.

The relationship between SC bioavailability and pI or local surface charge of mAbs was previously assessed using clinical or preclinical data in several reports[3, 10, 11]. For example, a weak inverse correlation was observed between SC bioavailability in monkeys and rats and local surface charge of six mAbs[11]. The local surface charge of the six mAbs was determined as the relative heparin-binding affinity of mAbs. Similarly, a weak inverse linear correlation (regression equation F = - 0.036 * pI + 1.0044, R^2^ = 0.054) was observed between human SC bioavailability and pI of 17 mAbs, where the pI was calculated based on the amino acid sequence of mAbs[10]. In contrast, no clear trend was observed between human SC bioavailability and pI of 14 mAbs, where pI data was collected from literature reports[3]. The poor correlation might be caused by the small sample size (N = 14) and cross-study variability in experimental pI measurement. In current study, sequence-based pI was calculated using the amino acid sequence of individual fusion protein and mAb while the 3D pI of Fv region was calculated by homology modeling[14], which eliminated cross-study variability in experimental pI measurement. Our analysis showed that 3D pI of Fv region was more correlated with SC bioavailability of mAbs (R^2^ = 0.174) than sequence-based pI (R ^2^= 0.092). The Fc domain of fusion proteins and mAbs usually shows negligible variability in electrostatic charge while the Fv regions of therapeutic proteins show significant diversity in electrostatic charge[14]. Thus, the 3D pI of Fv region is a more sensitive measure of potential charge-based interactions between therapeutic proteins and SC tissues than the sequence-based pI of the whole protein molecule. It was reported that mAbs with increased positive charge in Fv regions exhibited incrementally enhanced binding to epithelial cell surfaces and cellular uptake[12]. Furthermore, pharmacokinetic modeling of 19 SC administered mAbs revealed that electrostatic charge in the Fv regions substantially influenced SC absorption rate of mAbs[24].

Although the 3D pI of Fv region is more correlated with SC bioavailability than the sequence-based pI of whole mAb molecule, it is convenient to calculate sequence-based pI. We developed four multivariate regression models using either 3D pI of Fv region or sequence-based pI and compared the prediction performance of the four models (Table 1). It was not surprising that the linear regression model using 3D pI of Fv region generated the lowest GMFE of 1.127-fold, indicating this model should be selected when the pI of Fv region is available. The linear regression model using sequence-based pI generated a prediction accuracy comparable to that of paraboloid regression model using 3D pI of Fv region (PpwFE < 0.2 fold: 82% versus 79%; GMFE: 1.135-fold versus 1.133-fold). When the 3D pI of Fv region is not available, it is preferred to select the linear regression model using sequence-based pI for SC bioavailability prediction.

To the best of the author’s knowledge, no quantitative approach has been successfully developed and validated to predict human SC bioavailability in the absence of human SC pharmacokinetic data. A two-compartment model was developed to estimate clearance and SC bioavailability of mAbs using human SC pharmacokinetic data only[25]. The developed model was used to estimate SC bioavailability of 25 mAbs with linear pharmacokinetic profiles. The GMFE of the predictions was 1.195-fold and 60% of predictions were within 0.8-to 1.2-fold error. Although human SC pharmacokinetic data was needed in the two-compartment model, the prediction performance of this model was inferior to our multivariate regression models.

One limitation of our multivariate regression models is that human IV linear clearance is required as an independent variable. Human pharmacokinetic data of fusion proteins and mAbs may not be available at early stage of drug development, which precludes the application of these models to therapeutic proteins at preclinical stage. To address this knowledge gap, quantitative models using in vitro and preclinical data only are under development for human SC bioavailability predictions. The results will be reported separately. Another limitation of this study is the lack of model validation for Fc- and albumin-fusion proteins. Due to the small number of fusion protein (N = 20), model validation could not be conducted with an independent dataset. When the human SC bioavailability data of more fusion proteins is available, appropriate model validation is warranted.

## 5. Conclusions

In this study, an inverse linear correlation was observed between human SC bioavailability and human intravenous clearance or pI of 20 Fc-or albumin-fusion proteins and 98 monoclonal antibodies. Based on the observed relationships, the author developed multivariate regression models to predict human SC bioavailability of fusion proteins and mAbs. The prediction performance of the models was assessed using an independent test dataset. Two linear regression models based on the pI values of Fv region and whole molecule generated 73% and 82% of predictions within 0.8-to 1.2-fold deviations, respectively. Overall, this study demonstrated that clearance- and pI-based multivariate regression models could be used to predict human SC bioavailability of fusion proteins and mAbs.

