## Supplementary materials for "Predicting Human Bioavailability of Subcutaneously Administered Fusion Proteins and Monoclonal Antibodies"

Peng Zou[1*]

1. Quantitative Clinical Pharmacology, Daiichi Sankyo, Inc., 211 Mt. Airy Road, Basking Ridge, NJ 07920

*Corresponding author:

Peng Zou, Ph.D.

Quantitative Clinical Pharmacology

Daiichi Sankyo, Inc.

211 Mount Airy Road

Basking Ridge, NJ 07920

**Table S1. A summary of twenty Fc- or albumin-fusion proteins for regression analysis**

| Number | Fusion Proteins | F | pI& | CL (mL/day/kg) | References |
|---|---|---|---|---|---|
| 1 | Abatacept | 0.786 | 5.67 | 5.52 | 1 |
| 2 | Acazicolcept | 0.606 | 5.91 | 40.8 | 2 |
| 3 | Aflibercept | 0.549 | 8.29 | 12.9 | 3 |
| 4 | Alefacept | 0.63 | 6.11 | 6 | 4 |
| 5 | Asfotase alfa | 0.602 | 5.86 | 181 | 5 |
| 6 | Atacicept | 0.31 | 8.14 | 10.94 | 6 |
| 7 | Belatacept | 0.79 | 5.56 | 14.13 | 7 |
| 8 | Blisibimod | 0.412 | 6.49 | 16.66 | 8 |
| 9 | Coagulation Factor IX albumin fusion protein (IDELVION) | 0.55 | 5.47 | 20.6 | 9 |
| 10 | Dulaglutide | 0.65 | 5.45 | 14.61 | 10 |
| 11 | Efgartigimod | 0.5 | 7.25 | 37.03 | 11 |
| 12 | Etanercept | 0.58 | 7.2 | 26.2 | 12 |
| 13 | Fc-osteoprotegerin | 0.072 | 7.03 | 67.2 | 13 |
| 14 | Luspatercept | 0.76 | 5.4 | 4.78 | 14 |
| 15 | Rilonacept | 0.5 | 6.95 | 5.77 | 15 |
| 16 | Romiplostim | 0.35 | 8.3 | 157.9 | 16 |
| 17 | Sotatercept | 0.672 | 5.97 | 2.13 | 17-18 |
| 18 | UTTR1147A (IL-22Fc) | 0.68 | 7.81 | 24.6 | 19 |
| 19 | Inbakicept (IL-15Fc) | 0.0297 | 8.6 | 240 | 20 |
| 20 | Olamkicept | 0.48 | 6.31 | 56.2 | 21-22 |

Abbreviations: F, subcutaneous bioavailability; pI, isoelectric point; CL, clearance

&pI was calculated using Expasy (https://www.expasy.org/) based on amino acid sequence.

**Table S2. A summary of ninety-eight monoclonal antibodies with human subcutaneous bioavailability available**

| No. | mAbs | pI[1] | pI-3D[2] | BSA_LC_HC[2] | RP[2] | DM.HM[2] | Avg_HI[2] | F | CL (mL/day/kg) | TMDD-CL (mL/day/kg) | Refs |
|---|---|---|---|---|---|---|---|---|---|---|---|
| 1 | Abrilumab | 6.47 | 4.75 | 916.992 | 2.932 | 2.602 | 1.129 | 0.905 | 1.21 | 1.69 | 23 |
| 2 | Adalimumab | 8.36 | 8.05 | 733.166 | 2.274 | 0.584 | 1.167 | 0.64 | 3.94 | 5.14 | 24 |
| 3 | Aducanumab | 8.79 | 9.64 | 879.424 | 3.550 | 1.181 | 0.852 | 0.54 | 5.28 | 9.36 | 25 |
| 4 | Alemtuzumab | 8.68 | 9.5 | 859.502 | 2.320 | 0.595 | 1.146 | 0.32 | 14.6 | 21.6 | 26 |
| 5 | Alirocumab | 8.11 | 7.71 | 769.602 | 1.675 | 0.482 | 0.983 | 0.85 | 3.17 | 6.2 | 27 |
| 6 | Amivantamab | 8.36 | N.A. | N.A. | N.A. | N.A. | N.A. | 0.65 | 5.14 | N.A. | 28-29 |
| 7 | Anifrolumab | 7.92 | 6.93 | 739.084 | 2.619 | 1.220 | 0.787 | 0.87 | 3.34 | 6.36 | 30 |
| 8 | Anrukinzumab | 8.1 | 6.4 | 805.798 | 1.844 | 0.612 | 1.091 | 0.973 | 2.51 | N.A. | 31 |
| 9 | Atezolizumab | 8.25 | 7.23 | 798.469 | 1.226 | 1.623 | 1.300 | 0.71 | 3.14 | N.A. | 32 |
| 10 | Bapineuzumab | 8.25 | 7.9 | 780.693 | 1.848 | 1.012 | 0.702 | 0.7 | 2.43 | N.A. | 33 |
| 11 | Belimumab | 8.25 | 7.43 | 775.257 | 2.266 | 1.676 | 0.891 | 0.74 | 3.2 | N.A. | 34 |
| 12 | Benralizumab | 8.52 | 8.57 | 785.610 | 1.799 | 1.584 | 1.923 | 0.59 | 4.16 | 6.68 | 35 |
| 13 | Bimekizumab | 8.25 | 8.13 | 968.604 | 2.068 | 0.470 | 0.950 | 0.701 | 2.43 | 2.71 | 36 |
| 14 | Blosozumab | 6.19 | 5.49 | 833.656 | 2.750 | 2.037 | 1.052 | 0.69 | 3.02 | N.A. | 37 |
| 15 | Bococizumab | 8.58 | 9.67 | 841.368 | 2.412 | 0.770 | 1.589 | 0.45 | 4.74 | 9.81 | 38 |
| 16 | Brazikumab | 7.27 | 6.84 | 773.135 | 2.171 | 0.328 | 0.734 | 0.75 | N.A. | N.A. | 39 |
| 17 | Briakinumab | 8.74 | 9.42 | 876.464 | 3.889 | 0.774 | 1.165 | 0.52 | 6.3 | 15.2 | 40 |
| 18 | Brodalumab | 8.33 | 9.2 | 827.852 | 2.285 | 1.126 | 0.630 | 0.55 | 4.38 | 5.02 | 41 |
| 19 | Burosumab | 8.25 | 8.15 | 734.138 | 1.703 | 1.647 | 1.390 | 0.9 | 3.36 | 4.94 | 42 |
| 20 | Canakinumab | 8.24 | 7.78 | 787.628 | 2.163 | 1.598 | 0.750 | 0.7 | 2.77 | 2.46 | 43 |
| 21 | Casirivimab | 8.09 | N.A. | N.A. | N.A. | N.A. | N.A. | 0.718 | 2.41 | N.A. | 44 |
| 22 | CNTO 5825 | N.A. | N.A. | N.A. | N.A. | N.A. | N.A. | 0.75 | 2.55 | 4.78 | 45 |
| 23 | Concizumab | 6.28 | 6.09 | 854.237 | 2.903 | 1.648 | 0.636 | 0.472 | 13.68 | N.A. | 46 |
| 24 | Crenezumab | 6.67 | 7.18 | 654.866 | 0.976 | 0.954 | 0.626 | 0.662 | 2.3 | N.A. | 47 |
| 25 | Crovalimab | 8.45 | | | | | | 0.9 | 1.6 | N.A. | 48 |
| 26 | Daclizumab | 8.44 | 8.68 | 744.737 | 2.788 | 0.425 | 1.036 | 0.9 | 2.97 | N.A. | 49 |
| 27 | Daratumumab | 8.26 | 7.46 | 804.863 | 1.034 | 1.208 | 1.038 | 0.69 | 7.2 | N.A. | 50-51 |

| | | | | | | | | | | | |
|---|---|---|---|---|---|---|---|---|---|---|---|
| 28 | Denosumab | 8.23 | 8.73 | 784.709 | 1.332 | 1.304 | 1.006 | 0.62 | 1.45 | 4.63 | [52] |
| 29 | Domagrozumab | 8.36 | 8.09 | 780.991 | 1.477 | 0.970 | 1.024 | 0.62 | 2.68 | 3.58 | [53] |
| 30 | Donanemab | 8.44 | N.A. | N.A. | N.A. | N.A. | N.A. | 0.6 | 8.77 | 11.4 | [54] |
| 31 | Dupilumab | 6.86 | 7.81 | 943.370 | 1.818 | 1.619 | 1.149 | 0.64 | 4.18 | 7.14 | [55] |
| 32 | Efalizumab | 8.53 | 8.57 | 884.151 | 1.784 | 0.095 | 1.236 | 0.43 | 9.23 | 15.5 | [56] |
| 33 | Emicizumab | 6.48 | 5.3 | 817.351 | 3.323 | 0.541 | 0.841 | 0.868 | 3.26 | N.A. | [57] |
| 34 | Enokizumab | 7.9 | 6.86 | 1074.288 | 2.041 | 1.626 | 1.589 | 0.79 | 1.96 | N.A. | [58] |
| 35 | Eptinezumab | 7.89 | 6.23 | 592.639 | 1.250 | 1.001 | 0.602 | 0.717 | 2.25 | N.A. | [58] |
| 36 | Erenumab-aooe | 8.58 | 9.45 | 870.414 | 2.288 | 0.328 | 0.770 | 0.82 | 2.75 | N.A. | [59] |
| 37 | Etokimab | 8.78 | N.A. | N.A. | N.A. | N.A. | N.A. | 0.29 | 7.2 | N.A. | [58] |
| 38 | Etrolizumab | 8.25 | 7.87 | 772.990 | 2.212 | 0.974 | 1.398 | 0.504 | 4.33 | N.A. | [60] |
| 39 | Evinacumab | 6.86 | 6.67 | 914.881 | 1.867 | 1.202 | 0.972 | 0.71 | 2.53 | 3.91 | [61] |
| 40 | Evolocumab | 8.24 | 7.49 | 666.156 | 2.980 | 0.825 | 0.708 | 0.72 | 4.1 | N.A. | [62] |
| 41 | Fremanezumab | 7.63 | 7.56 | 796.598 | 1.640 | 0.988 | 0.633 | 0.66 | 1.29 | N.A. | [63] |
| 42 | Fulranumab | 7.64 | 8.05 | 879.847 | 2.300 | 3.105 | 0.710 | 0.83 | 3.13 | N.A. | [64] |
| 43 | Gevokizumab | 8.08 | 9.22 | 871.008 | 2.289 | 0.689 | 0.213 | 0.69 | 2.5 | N.A. | [65] |
| 44 | Golimumab | 8.54 | 8.92 | 872.758 | 1.759 | 1.606 | 0.852 | 0.53 | 4.9 | 6.7 | [66] |
| 45 | Guselkumab | 8.45 | 8.72 | 763.839 | 1.448 | 1.061 | 0.303 | 0.49 | 4.61 | 6.03 | [67-68] |
| 46 | Ianalumab | 8.55 | 8.98 | 797.381 | 1.043 | 0.794 | 0.955 | 0.47 | 8.7 | 12.36 | [69] |
| 47 | Imdevimab | 8.44 | N.A. | N.A. | N.A. | N.A. | N.A. | 0.717 | 2.97 | N.A. | [44] |
| 48 | Imsidolimab | 8.36 | N.A. | N.A. | N.A. | N.A. | N.A. | 0.9 | 2.96 | N.A. | [70] |
| 49 | Inebilizumab | 6.96 | 5.88 | 834.815 | 3.139 | 2.203 | 0.846 | 0.81 | 2.69 | N.A. | [71] |
| 50 | Infliximab | 7.33 | 5.48 | 755.504 | 1.794 | 2.576 | 0.686 | 0.791 | 5.07 | N.A. | [72] |
| 51 | Iscalimab | 8.11 | 7.85 | 677.161 | 1.934 | 0.956 | 0.593 | 0.661 | 4.62 | N.A. | [73] |
| 52 | Ixekizumab | 7.92 | 8.84 | 820.732 | 1.567 | 1.181 | 1.120 | 0.705 | 5.57 | 10.01 | [74] |
| 53 | Lebrikizumab | 6.43 | 4.88 | 805.156 | 1.340 | 1.029 | 0.452 | 0.856 | 1.97 | N.A. | [75] |
| 54 | Lecanemab | 8.45 | N.A. | N.A. | N.A. | N.A. | N.A. | 0.497 | 5.38 | 9.69 | [76-77] |
| 55 | Leronlimab | 7.3 | N.A. | N.A. | N.A. | N.A. | N.A. | 0.5 | 21.8 | N.A. | [78] |
| 56 | Ligelizumab | 7.32 | 5.81 | 826.002 | 1.286 | 0.965 | 1.028 | 0.78 | 7.42 | 14.1 | [79] |
| 57 | Lirentelimab | 8.11 | N.A. | N.A. | N.A. | N.A. | N.A. | 0.63 | N.A. | N.A. | [80] |

| | | | | | | | | | | | |
|---|---|---|---|---|---|---|---|---|---|---|---|
| 58 | Lutikizumab | 8.77 | N.A. | N.A. | N.A. | N.A. | N.A. | 0.46 | 4.04 | 3.97 | [81] |
| 59 | Marstacimab | 7.93 | 5.58 | 637.048 | 1.725 | 2.275 | 0.533 | 0.373 | 10.49 | 19.72 | [82] |
| 60 | Mepolizumab | 8.37 | 8.12 | 835.589 | 1.871 | 1.210 | 0.907 | 0.8 | 2.29 | 3.29 | [83] |
| 61 | Mosunetuzumab | 8.49 | 8.25 | 774.309 | 2.800 | 0.362 | 0.373 | 0.75 | 8.34 | 15.4 | [84-85] |
| 62 | Natalizumab | 7.9 | 8.68 | 835.939 | 1.904 | 0.257 | 1.228 | 0.66 | 3.5 | N.A. | [86] |
| 63 | Ofatumumab | 8.36 | 8.05 | 791.486 | 1.782 | 1.606 | 0.493 | 0.685 | 4.86 | N.A. | [87] |
| 64 | Olokizumab | 6.27 | 5.06 | 780.104 | 1.300 | 0.531 | 0.467 | 0.842 | 1.99 | 2.45 | [58, 88] |
| 65 | Omalizumab | 7.35 | 5.48 | 877.553 | 1.747 | 1.031 | 0.941 | 0.62 | 2.59 | N.A. | [89] |
| 66 | Opicinumab | 7.29 | 5.21 | 811.523 | 1.893 | 0.995 | 0.545 | 0.65 | 3.57 | 5.86 | [90] |
| 67 | Otilimab | 8.61 | N.A. | N.A. | N.A. | N.A. | N.A. | 0.35 | 13.3 | N.A. | [91] |
| 68 | Pateclizumab | 8.53 | 8.84 | 796.923 | 1.542 | 0.539 | 0.903 | 0.43 | 7.03 | N.A. | [92] |
| 69 | Pembrolizumab | 7.63 | 8.37 | 720.807 | 1.631 | 0.872 | 1.024 | 0.64 | 3.06 | 4.72 | [93] |
| 70 | Pertuzumab | 8.35 | 8.41 | 806.191 | 1.247 | 1.316 | 0.566 | 0.7 | 2.55 | N.A. | [94] |
| 71 | PF-04236921 | N.A. | N.A. | N.A. | N.A. | N.A. | N.A. | 1 | 1.66 | N.A. | [95] |
| 72 | Prezalumab | 7.26 | 7.69 | 724.787 | 1.275 | 0.638 | 1.019 | 0.57 | 8.14 | N.A. | [96] |
| 73 | Pozelimab | 7.23 | N.A. | N.A. | N.A. | N.A. | N.A. | 0.7 | N.A. | N.A. | [97] |
| 74 | Ralpancizumab | 8.5 | 9.58 | 777.790 | 2.639 | 0.712 | 1.277 | 0.62 | 4.93 | 3.58 | [98] |
| 75 | Ravulizumab | 6.32 | 6.15 | 896.413 | 2.465 | 1.320 | 1.223 | 0.79 | 1.14 | N.A. | [99] |
| 76 | Remtolumab | 6.85 | N.A. | N.A. | N.A. | N.A. | N.A. | 0.48 | 4.55 | 9.76 | [100] |
| 77 | Reslizumab | 7.1 | 6.79 | 814.976 | 1.477 | 1.284 | 0.530 | 0.67 | 2.64 | 3.26 | [101] |
| 78 | Risankizumab | 8.36 | 8.88 | 830.016 | 1.957 | 0.523 | 0.892 | 0.89 | 3.43 | 4.64 | [102] |
| 79 | Rituximab | 8.66 | 9.33 | 848.375 | 1.868 | 0.487 | 0.801 | 0.58 | 4.79 | N.A. | [103] |
| 80 | Romosozumab | 6.67 | 5.19 | 837.445 | 2.938 | 1.451 | 1.515 | 0.81 | 5.04 | 8.69 | [104] |
| 81 | Rontalizumab | 8.25 | 7.52 | 816.046 | 1.796 | 1.638 | 0.735 | 0.44 | 2.88 | N.A. | [105] |
| 82 | Rosnilimab | 8.09 | N.A. | N.A. | N.A. | N.A. | N.A. | 0.8 | N.A. | N.A. | [106] |
| 83 | Sarilumab | 8.26 | 7.52 | 762.812 | 2.116 | 1.983 | 1.046 | 0.8 | 2.97 | N.A. | [107] |
| 84 | Sasanlimab | 7.63 | N.A. | N.A. | N.A. | N.A. | N.A. | 0.6 | 4.75 | 4.93 | [108] |
| 85 | Satralizumab | 5.92 | 5.1 | 778.728 | 3.840 | 2.166 | 0.389 | 0.85 | 2.4 | 12.3 | [109] |
| 86 | Secukinumab | 8.09 | 7.58 | 827.869 | 1.651 | 0.827 | 0.607 | 0.73 | 2.09 | N.A. | [110] |
| 87 | Sifalimumab | 8.61 | 9.26 | 698.814 | 2.152 | 1.114 | 0.796 | 0.75 | 2.32 | N.A. | [111-112] |

| | | | | | | | | | | | |
|---|---|---|---|---|---|---|---|---|---|---|---|
| 88 | Sirukumab | 7.92 | 5.39 | 745.948 | 1.306 | 1.487 | 1.238 | 0.87 | 3.8 | 6.1 | [113] |
| 89 | Tabalumab | 8.1 | 8.97 | 913.493 | 1.465 | 0.343 | 0.473 | 0.62 | 3.94 | 5.29 | [114] |
| 90 | Tanezumab | 7.88 | 8.1 | 917.691 | 1.298 | 0.578 | 0.903 | 0.69 | 2.41 | N.A. | [115] |
| 91 | Teclistamab | 8.11 | N.A. | N.A. | N.A. | N.A. | N.A. | 0.69 | 4.43 | N.A. | [116] |
| 92 | Tezepelumab | 7.33 | 6.61 | 771.484 | 2.041 | 0.893 | 0.971 | 0.81 | 2.3 | N.A. | [117] |
| 93 | Tildrakizumab | 8.35 | 8.29 | 706.401 | 1.531 | 0.840 | 1.823 | 0.73 | 2.11 | 2.52 | [118] |
| 94 | Tocilizumab | 8.62 | 9.14 | 750.796 | 1.314 | 1.160 | 1.240 | 0.795 | 4.43 | 5.14 | [119-120] |
| 95 | Tralokinumab | 6.88 | 5.3 | 849.301 | 1.688 | 0.783 | 0.802 | 0.76 | 2.13 | N.A. | [121] |
| 96 | Trastuzumab | 8.45 | 8.41 | 735.698 | 1.586 | 0.485 | 0.975 | 0.77 | 1.59 | 5.01 | [122] |
| 97 | Ustekinumab | 8.53 | 8.5 | 797.384 | 1.611 | 0.495 | 1.156 | 0.572 | 2.3 | N.A. | [123] |
| 98 | Vedolizumab | 8.09 | 7.39 | 916.048 | 1.570 | 0.751 | 1.205 | 0.75 | 2.24 | N.A. | [124] |

Abbreviations: mAbs = monoclonal antibodies; pI = isoelectric point; pI-3D = structure-based isoelectric point of variable regions; BSA_LC_HC = surface area buried between variable regions of light and heavy chains; RP = ratio of charged to hydrophobic surface patches; DM.HM = Ratio of dipole and hydrophobic moments; Avg_HI = average hydrophobic imbalance; F = subcutaneous bioavailability; CL = clearance; TMDD_CL = target-mediated drug disposition clearance or nonlinear clearance.

[1] pI was calculated using Expasy (https://www.expasy.org/) based on amino acid sequence.

[2] pI-3D, BSA_LC_HC, RP, DM.HM, and Avg_HI were calculated using both the energy minimized models and conformer ensembles generated using LowModeMD as implemented in Molecular Operating Environment (MOE) platform [125].

**Figure S1. Correlation analysis between intravenous linear clearance and (A) sequence-based pI or (B) 3D pI of Fv region**

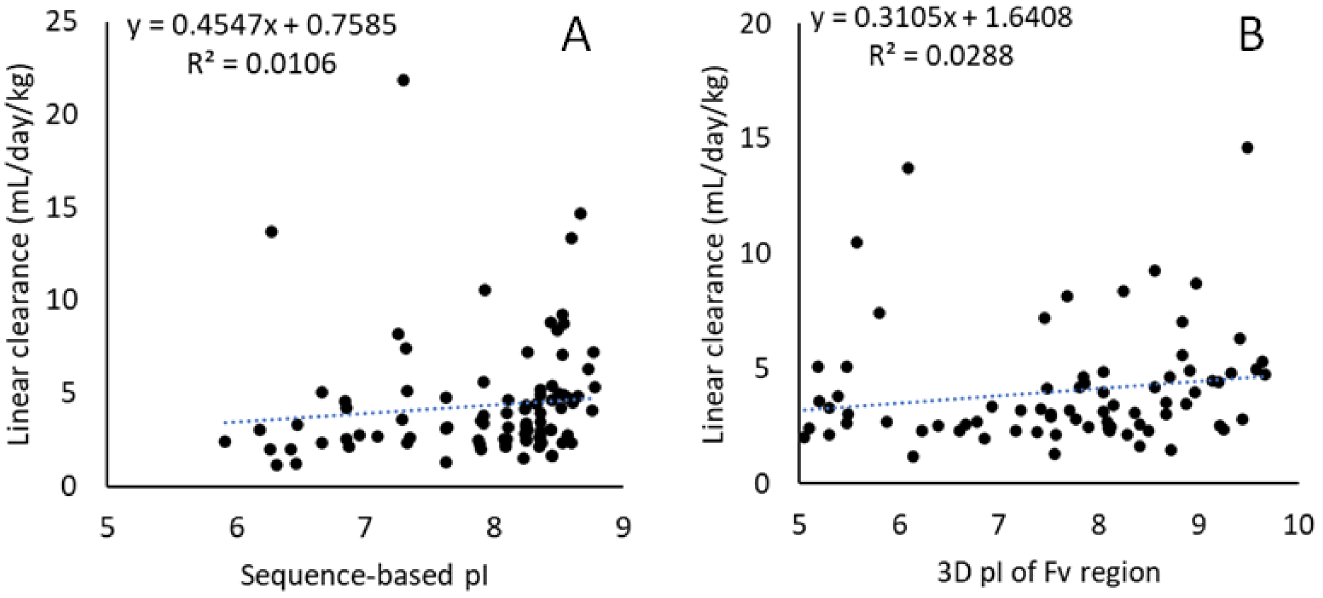
